# Multimodal EEG-fNIRS Fusion for Passive BCI-based Depressive State Classification

**DOI:** 10.64898/2026.04.07.716028

**Authors:** Riki Sakurai, Simon Kojima, Mihoko Otake-Matsuura, Shin’ichiro Kanoh, Tomasz M. Rutkowski

**Author notes:** Research and data analysis were performed while the author held a concurrent affiliation with the RIKEN Center for Advanced Intelligence Project (AIP), Tokyo, Japan.

## Abstract

Traditional psychiatric assessments for depression are often hindered by subjective bias and patient recall inaccuracy. This paper presents a multimodal passive Brain-Computer Interface (pBCI) designed for the objective screening of depressive traits through the end-to-end decoding of neural dynamics. We implemented a hybrid EEG-fNIRS framework to capture synchronized electro-hemodynamic responses during an emotional working memory (EWM) task. To classify sub-clinical depressive tendencies based on BDI-II scores, we utilized SincShallowNet, a deep learning architecture optimized for raw signal processing via learnable Sinc-filters. Our results demonstrate that the pBCI achieves peak performance in the auditory modality, with the integration of EEG and low-pass filtered fNIRS (0.15 Hz) yielding a balanced accuracy of 90.9% and an F1-score of 0.867. By isolating purely endogenous neural markers during the EWM maintenance phase, the system provides a robust “silent observer” for mental state monitoring. These findings validate the potential of multimodal pBCIs as high-precision, data-driven tools for early-stage depression screening, offering a scalable alternative to traditional clinical interviews and a foundation for longitudinal mental health monitoring.

## I. Introduction

A passive brain-computer interface (pBCI) [1] is a system that monitors and interprets spontaneous brain activity occurring without the user’s explicit intent to control a device. Unlike Active BCIs, which are oriented toward command generation – such as imagining hand movements to control a cursor or mentalizing musical scores to play chords [2] – or Reactive BCIs, which rely on external stimuli like flashing visual patterns or sequences of auditory/tactile stimulations to evoke specific neural responses [3], a pBCI acts as a “silent observer” [4]. It implicitly extracts information regarding the user’s cognitive or affective state–such as mental workload, stress, fatigue, or mood–while the user remains engaged in a primary task.

In response to the surging global demand for mental healthcare, pBCI have emerged as a transformative tool for the objective assessment of Major Depressive Disorder (MDD) [5]. Unlike active BCIs that require conscious task execution, pBCIs seamlessly monitor the user’s ongoing neurophysiological state without interrupting their daily activities. Traditional diagnostic frameworks remain heavily reliant on clinical interviews and longitudinal self-reports, such as the Beck Depression Inventory II (BDI-II) [6]. While foundational, these methods suffer from:

- **Subjective Bias:** Patient reports are often clouded by memory recall issues or the psychological state itself.
- **Clinical Subjectivity:** Diagnoses can vary based on a clinician’s individual empirical judgment.
- **Snapshot Constraints:** Interviews provide only a “point-in-time” assessment rather than capturing the fluctuating nature of depressive symptoms.

To overcome these hurdles, the integration of Electroencephalography (EEG) and functional Near-Infrared Spectroscopy (fNIRS) within a pBCI framework provides a “gold standard” for quantitative mental health monitoring [7]. The integration of these two modalities allows for a synergistic evaluation that is far less susceptible to conscious bias than traditional assessments. By leveraging pBCIs, clinicians can move toward a predictive model of care–detecting subtle shifts in neural oscillations and blood oxygenation that precede a clinical relapse. This objective data stream transforms depression management from a reactive “crisis-response” model [8] into a proactive, data-driven monitoring system.

Early intervention is paramount in altering the trajectory of MDD. The pBCI systems offer a unique advantage here, as they can identify latent neurophysiological markers – such as deficits in emotional working memory (EWM) – that often precede a patient’s conscious awareness of worsening symptoms. Unlike active diagnostics, this pBCI framework operates by monitoring the brain’s “background” processing of affective information. Since impaired EWM is a core cognitive hallmark of depression, the system analyzes neural efficiency during tasks that require the retention of emotional stimuli. To capture a high-fidelity representation of these mental states, we implemented a dual-modality approach:

- **Synchronized Acquisition:** Participants engaged in emotional short-term memory tasks while EEG (capturing rapid inhibitory oscillations) and fNIRS (measuring prefrontal oxygenation levels) were recorded simultaneously.
- **Ground Truth Mapping:** These objective neural signals were labeled against BDI-II [6] scores, allowing the system to correlate specific neuro-hemodynamic patterns with established clinical severity.

The core of this screening system is SincShallowNet [9]– [11], a deep learning architecture optimized for pBCI applications. By leveraging the strengths of both SincNet [9] (for interpretable frequency filtering) and ShallowConvNet [10] (for spatial feature extraction), the model effectively decodes complex neural dynamics. Our objective is to move beyond subjective reporting by using SincShallowNet [11] to estimate depression levels automatically, providing a continuous, objective monitoring tool that detects “silent” shifts in a user’s psychological state.

## II. Methods

The study cohort comprised eleven healthy university students (5 female, 6 male) with a mean age of 22.27 *±* 1.21 years. Experimental sessions were hosted at the RIKEN Center for Advanced Intelligence Project (AIP) in Tokyo, Japan. All procedures were conducted in strict accordance with the Declaration of Helsinki and received formal approval from the RIKEN Ethical Committee for Experiments with Human Subjects (Permission No. Wako3 30-28(4)).

To facilitate a comprehensive pBCI framework, EEG and fNIRS data were recorded simultaneously. This multimodal approach was designed to capture both the rapid electrical oscillations and the slower hemodynamic responses associated with affective processing.

### A. Affective Ground Truth Assessment

A critical component of the passive screening framework is the establishment of objective labels for the training of machine-learning models. Prior to neural recording, participants’ baseline affective states were quantified using the BDI-II [6].

- **Instrument Modification:** A 20 −item version of the BDI-II was implemented, omitting the item related to sexual interest to ensure participant comfort and uphold ethical privacy standards.
- **Labeling Strategy:** These scores served as the supervisory signal (ground truth) for the pBCI, allowing the system to learn the mapping between spontaneous neural dynamics and different levels of depressive symptoms.

### B. Experimental Protocol

To elicit neural biomarkers associated with MDD [5], we implemented a multimodal pBCI experimental framework. The paradigm was designed to probe the neurophysiological dynamics of EWM, as cognitive deficits in this domain are recognized as key indicators of depressive tendencies.

#### 1) Stimulus Selection and Presentation

Participants were presented with emotional expressions – sourced from the “Mind Reading” database [12] – delivered through two distinct modalities:

- **Visual Stimuli:** Short video clips of facial expressions.
- **Auditory Stimuli:** Spoken affective utterances.

#### 2) Experimental Structure and Session Design

The experiment was partitioned into six distinct sessions, following a 2 *×* 3 design:

1. **Modality:** Visual or Auditory.
2. **Affective Intensity:** Easy, Middle, and Hard subsets (derived from |*V* | and |*A*| ratings as detailed in Table I).

**TABLE 1:**
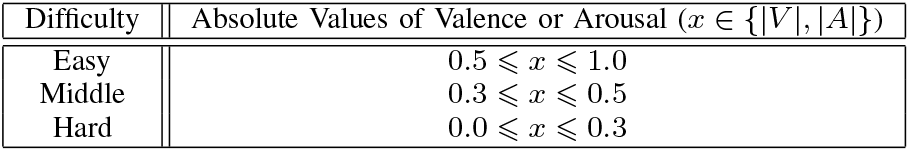
Absolute value range of valence (|*V*|) and arousal (|*A*|) ratings for affective auditory and visual stimuli sourced from the “Mind Reading” database [12]. These values delineate the emotional target subsets (Easy, Middle, and Hard) utilized as labels for passive BCI training.

Each session comprised ten trials. This structured approach allows the pBCI to be trained on varying levels of affective complexity, mirroring the subtle emotional processing nuances required for accurate depression prediction.

#### 3) Artifact Mitigation and pBCI Signal Integrity

To ensure the high signal-to-noise ratio (SNR) required for deep learning architectures like SincShallowNet [11], the protocol incorporated a delayed-response mechanism. Participants were instructed to withhold behavioral responses – provided via the hardware interface – until the completion of the affective stimulus. By isolating the cognitive “encoding” phase from the motor “response” phase, we successfully minimized electromyographic (EMG) interference and motion artifacts, preserving the integrity of the underlying EEG and fNIRS neural dynamics.

#### 4) Stimulus Selection and Affective Labeling for pBCI Training

To facilitate the supervised training of the classification model, stimuli were strategically selected to produce distinct neural signatures corresponding to varying affective intensities. Stimulus parameters were derived from the NRC VAD Lexicon [13], which provides normative valence (*V*), and arousal (*A*) scores on a scale from −1 to 1. We established three difficulty levels (Easy, Middle, and Hard) by thresholding the valence and arousal ranges (see Table I). This stratification is based on the hypothesis that higher affective intensity – represented by larger absolute values |*V*| and |*A*| – elicits more robust, easily discriminable neural responses, whereas values near zero represent subtle emotional states that increase cognitive demand for the subjects and produce more complex, overlapping features for the hybrid EEG-fNIRS classification boundaries.

The selection process for data labeling and trial construction was designed to maximize class discriminability for the machine-learning pipeline:

- **Ground-Truth Target Selection:** A target emotional word was randomly sampled from a pool meeting the specific |*V*| and |*A*| criteria for the assigned difficulty level. This word serves as the primary class label for the trial.
- **Contrastive Distractor Selection:** To ensure the model learns to distinguish between opposing affective states – essential for identifying EWM modulations associated with normal versus depressive mental states – a “non-target” word was selected with a valence sign opposite to that of the target.
- **Multimodal Training Samples:** Three media files were retrieved from the database to populate the trial: two distinct samples of the target emotion (one for the encoding phase and one for the classification-target phase) and one non-target sample. This ensures the pBCI model learns generalized depression-related features rather than specific media artifacts.

### C. EEG and fNIRS Data Acquisition Protocol

The experimental procedure was designed to isolate specific neural markers of EWM for subsequent machine-learning classification. Each trial followed a synchronized temporal sequence:

- **Target Phase (Encoding):** Participants were presented with a five-second “target stimulus” (affective video or audio). This phase serves as the initial neural encoding period, during which the pBCI system captures the neurophysiological responses to specific valence and arousal intensities as markers of the subject’s underlying depressive state.
- **Maintenance Phase (Working Memory):** A blank screen or neutral audio period followed, during which subjects maintained the emotional state in memory. For pBCI applications, this phase is critical as it provides a window of purely endogenous neural activity, free from primary sensory stimulation, for feature extraction.
- **Test Phase (Retrieval and Discrimination):** Two new affective stimuli were presented simultaneously to probe the accuracy of the stored emotional representation. One matched the target emotion, while the other served as a contrastive distractor.
  - *Video Modality:* Synchronized side-by-side play-back.
  - *Audio Modality:* Dichotic presentation with a one-second onset delay in one channel to decouple the recognition of speech from the onset of neural processing.
- **Behavioral Ground Truth (Response):** Participants performed a forced-choice task using the hardware interface (Figure 1). This task serves as a structured mental activity, consistent with dual-task pBCI paradigms, ensuring sustained cognitive engagement without contributing motor-related features to the classification pipeline, while the system passively decodes the underlying neural signatures.
- **Artifact Management (Self-Paced Protocol):** To maintain high signal quality and reduce artifacts caused by fatigue or eye-blink frequency, a self-pacing mechanism was utilized. Subjects initiated the next trial via the central blue button only when in a state of attentional readiness.

**Fig. 1:**
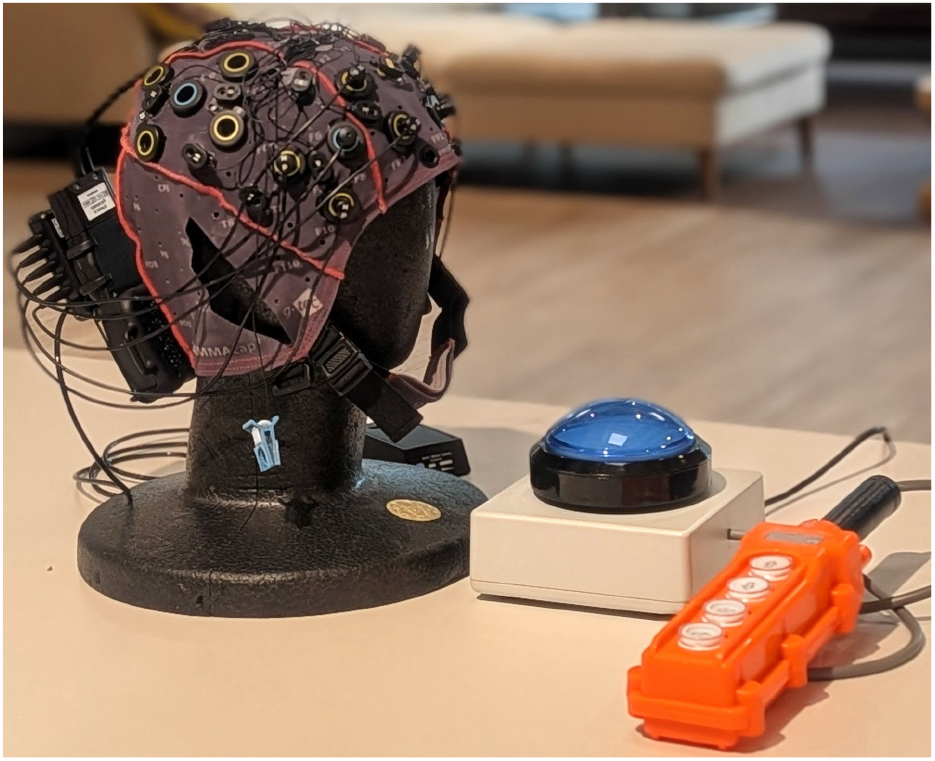
Experimental setup for multimodal pBCI data acquisition. During experiments, the participant is equipped with the g.NAUTILUS fNIRS system (g.tec medical engineering GmbH, Austria). The hardware interface features an orange response panel for target stimulus selection and a central blue button for self-paced trial progression. This configuration enables the synchronized recording of neural dynamics and behavioral markers during the affective stimulus paradigm, providing the labeled dataset required to train machine-learning models for the classification of normal versus depressive mental states.

To ensure the millisecond-precision required for supervised machine learning, all neural signals (EEG/fNIRS) and experimental event markers were unified and timestamped via the Lab Streaming Layer (LSL) protocol [14]. This creates the time-locked dataset illustrated in Figure 2, enabling the training of deep-learning models on specific cognitive epochs.

**Fig. 2:**
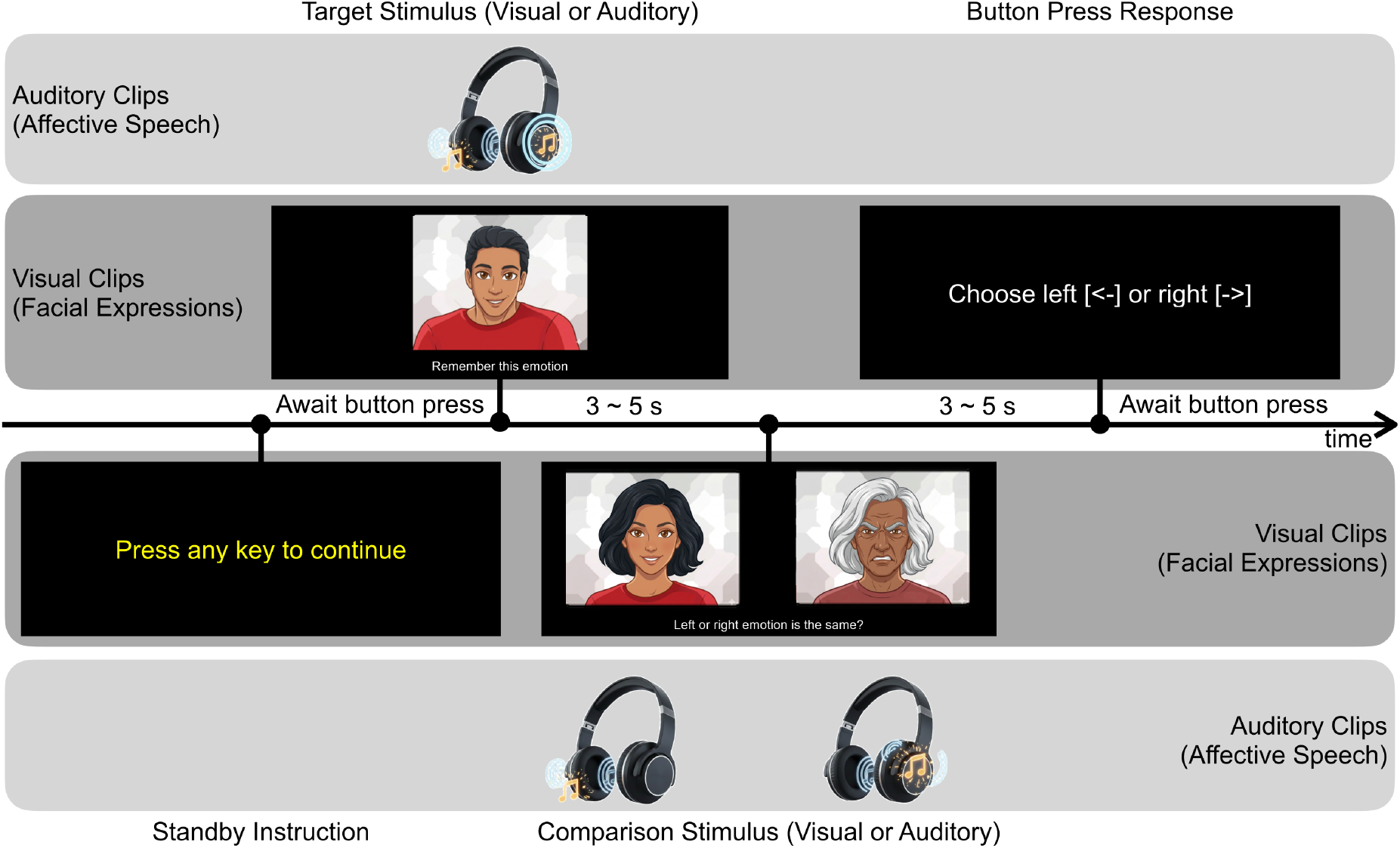
Experimental paradigm and stimulus presentation. Affective stimuli, consisting of Auditory Clips (affective speech) or Visual Clips (facial expressions), were presented in two phases. Note: while real video stimuli were used during the experiment, AI-generated drawings are used in this figure for privacy to mask actor identities while preserving the experimental affect. Initial encoding involved a single Target Stimulus for 3–5s, followed by a retrieval phase after a 3–5s delay. In the retrieval phase, stimuli were presented simultaneously (dichotically for audio; side-by-side for video) to evaluate emotional working memory. Subjects indicated if the emotion was the same via a Button Press Response.

### D. Multimodal pBCI Data Acquisition and Preprocessing

The system utilized a hybrid EEG–fNIRS g.NAUTILUS fNIRS platform (g.tec medical engineering GmbH, Austria) to capture the concurrent electro-hemodynamic signatures of affective processing. The spatial distribution of this multimodal sensor montage is illustrated in Figure 3.

**Fig. 3:**
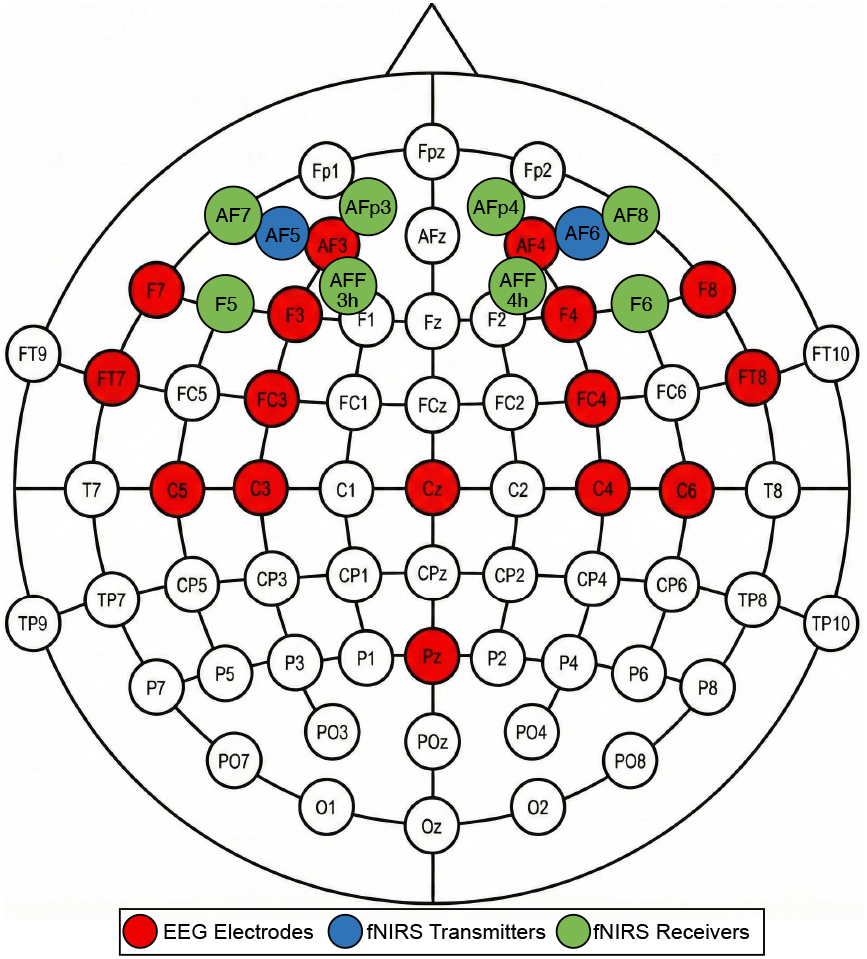
Hybrid EEG-fNIRS sensor topography. Schematic representation of the optode and electrode montage for the g.NAUTILUS fNIRS wireless system (g.tec medical engineering GmbH, Austria). EEG electrode sites are denoted by red markers, while fNIRS light sources (transmitters) and detectors (receivers) are represented by green and blue markers, respectively, illustrating the integrated spatial distribution across the scalp.

1. *EEG Configuration:* Sixteen active EEG electrodes were positioned at AF3, AF4, F7, F8, F3, F4, FC3, FC4, C5, FT7, Cz, FT8, C6, C3, C4, and Pz. This configuration focuses on the prefrontal and frontal cortices, regions highly associated with emotional regulation and working memory. Data were hardware-filtered (1 Hz high-pass; 50 Hz notch) to preserve signal integrity for real-time pBCI monitoring.
2. *fNIRS Configuration:* Cortical hemodynamic activity was monitored via an eight-channel subsystem integrated into the hybrid cap. The optode arrangement – comprising eight sources (AFp4, AF8, F6, AFF4h, AF7, AFp3, AFF3h, F5) and two detectors (AF6, AF5) – was designed to overlap with the EEG sites. A fixed source-detector separation of 32 mm was utilized to ensure sufficient cortical penetration, allowing the pBCI to track oxygenation changes (*HbO* and *HbR*) alongside electrical oscillations.
3. *pBCI Preprocessing Pipeline:* Multimodal signals were synchronously digitized at 250 Hz. To prepare the raw data for end-to-end classification, three-second epochs were extracted, precisely time-locked to the stimulus comparison onset via Lab Streaming Layer (LSL) software triggers. This windowing strategy isolates the peak cognitive load period, during which the pBCI decodes the latent neural signatures of the comparison task. The signals underwent the following transformations:

- **EEG Preprocessing:** A fourth-order Butterworth band-pass filter (3–45 Hz) was applied to isolate *θ, α, β*, and low −*γ* bands, which are primary pBCI indicators of cognitive workload and affective state.
- **fNIRS Preprocessing:** Raw optical intensities (760 nm and 850 nm) were converted to optical density changes. To determine the optimal frequency range for characterizing “passive” mood indicators, we evaluated three distinct low-pass filtering configurations: (i) 0.15 Hz to isolate pure hemodynamic neural activity; (ii) 0.6 Hz to incorporate respiratory fluctuations; and (iii) 2.0 Hz to include cardiac frequencies. This comparative approach allowed us to empirically validate which spectral bandwidth maximizes feature discriminability for the machine-learning architecture, as detailed in the subsequent results Section III.

### E. Ground Truth Labeling for Screening Scenarios

To train the pBCI as a screening tool, we categorized participants into two groups based on the BDI-II [6]. A cutoff score of 7 was adopted to define the classification targets:

- *Low-Score (LS) Group (*≤ 6): Assigned binary label 0.
- *High-Score (HS) Group (*≥ 7): Assigned binary label 1.

While the HS group represents “minimal depression” clinically [6], in a pBCI context, these labels represent sub-clinical neurophysiological tendencies. The objective of the machine-learning pipeline is to identify whether the “silent observer” can detect these subtle differences during the EWM task, effectively serving as an early-warning screening mechanism.

### F. End-to-End Multimodal Classification: SincShallowNet

To decode depressive states from the integrated 32-channel dataset (16 EEG + 16 fNIRS), we employed SincShallowNet [9], implemented via the Braindecode framework [11]. This architecture is specifically suited for pBCIs because it learns optimal filter banks directly from raw, noisy data. The model’s pipeline consists of three primary functional stages:

1. **Parameterized Temporal Filtering:** The first layer utilizes Sinc-functions to perform band-pass filtering. By learning only the cutoff frequencies (*f*_1_, *f*_2_), the model extracts physically interpretable features – such as frontal alpha asymmetry or hemodynamic slopes – while minimizing the risk of overfitting common in small-sample BCI datasets.
2. **Multimodal Spatial Integration:** A spatial depthwise convolution integrates features across all 32 channels. This allows the network to learn the cross-modal correlations between rapid electrical spikes and gradual oxygenation changes, providing a holistic view of the participant’s mental state.
3. **Log-Variance Feature Transformation:** The architecture applies a square activation followed by mean pooling and a logarithmic transformation. This approximates the log-power of the neural signal, a standard feature in BCI literature for identifying state-dependent changes in brain activity.

The core of our classification pipeline is the SincShallowNet architecture [9], implemented via the Braindecode framework [10], [11]. This model was selected for its ability to perform end-to-end decoding of raw neurophysiological signals, which is essential for pBCI systems aiming to minimize human-in-the-loop intervention. The SincShallowNet integrates the hierarchical spatial-temporal processing of the ShallowConvNet [10] with parameterized Sinc-filters [9]. Unlike standard convolutional layers that learn every weight in a kernel, the Sinc-layer only learns the low (*f*_1_) and high (*f*_2_) cutoff frequencies. This provides several advantages for our passive depression-screening task:

- *Physiological Interpretability:* The model automatically converges on the most relevant spectral bands (e.g., *α* or *θ* bands in EEG, or specific hemodynamic frequencies in fNIRS) for distinguishing depressive states.
- *Reduced Overfitting:* By significantly reducing the number of learnable parameters in the first layer, the architecture remains robust even with the relatively small datasets typical of clinical pBCI studies.
- *Multimodal Fusion:* The depthwise spatial convolution layer that follows the Sinc-filters effectively fuses the electrical (EEG) and hemodynamic (fNIRS) features into a unified latent representation of the user’s affective state.

The machine-learning pipeline was implemented using a Leave-One-Subject-Out (LOSO) cross-validation protocol to simulate real-world pBCI deployment, where the system must classify data from an unseen user. The evaluation was iterated eleven times – corresponding to the total number of participants – ensuring that each subject’s data served as a completely independent test set exactly once. This approach rigorously assesses the model’s ability to generalize across diverse neurophysiological profiles without subject-specific calibration.

## III. Results

We evaluated the decoding efficacy of the SincShallowNet architecture across the multimodal EEG-fNIRS dataset to assess its viability as a passive screening tool using a subject-independent LOSO cross-validation protocol. The performance metrics – summarized in Figure 4 (confusion matrices) and Table II (balanced accuracy, F1-score, precision, and recall) – demonstrate that the pBCI effectively identifies depressive indicators through spontaneous neural responses.

**Fig. 4:**
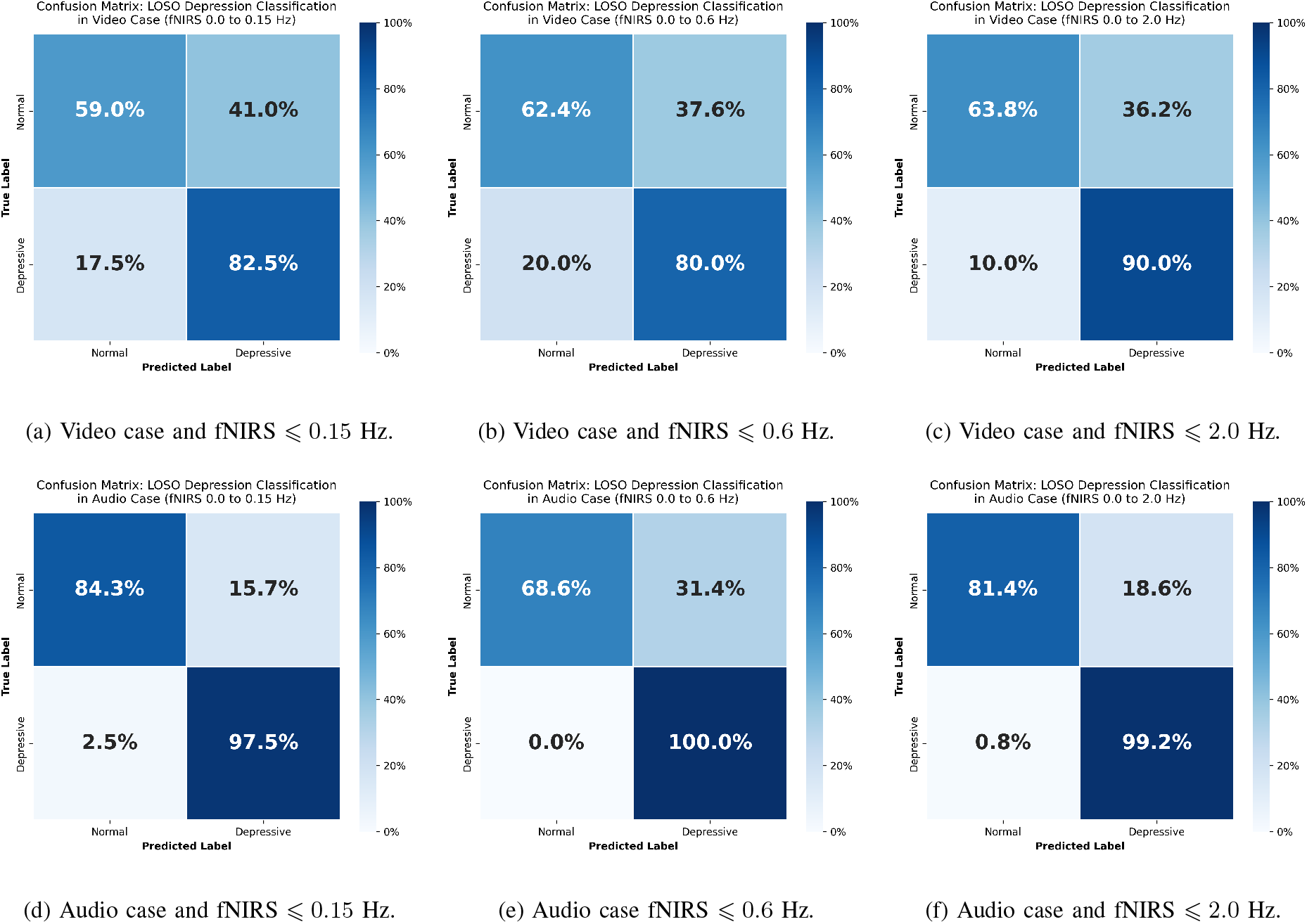
Confusion matrices for Leave-One-Subject-Out (LOSO) cross-validation. Panels (a)–(c) represent the video modality, and panels (d)–(f) represent the audio modality. Columns correspond to the three fNIRS low-pass filtering tiers: 0.15 Hz (left), 0.6 Hz (center), and 2.0 Hz (right), illustrating the impact of physiological frequency inclusion on classification accuracy.

**TABLE II.** Classification performance across different experimental conditions. Metrics include balanced accuracy, F1 score, precision, and recall for depression detection, with the best-performing results indicated in bold.

| Stimulus Setting | fNIRS Low-pass Filter | Balanced Accuracy | F1 Score | Precision | Recall |
| --- | --- | --- | --- | --- | --- |
| Video | fNIRS ≤ 0.15 Hz | 70.8% | 0.649 | 0.535 | 0.825 |
| Video | fNIRS ≤ 0.60 Hz | 71.2% | 0.651 | 0.549 | 0.800 |
| Video | fNIRS ≤ 2.00 Hz | 76.9% | 0.711 | 0.587 | 0.900 |
| Audio | fNIRS ≤ 0.15 Hz | **90.9%** | **0.867** | **0.780** | 0.975 |
| Audio | fNIRS ≤ 0.60 Hz | 84.3% | 0.784 | 0.645 | **1.000** |
| Audio | fNIRS ≤ 2.00 Hz | 90.3% | 0.856 | 0.753 | 0.992 |

The results highlight a superior predictive power within the auditory affective speech condition. Within this paradigm, the fNIRS low-pass filtering setting of 0.15 Hz – optimized to isolate pure hemodynamic neural activity – yielded the highest discriminability. This suggests that the pBCI most accurately captures depressive biomarkers when focused on slow-wave hemodynamic shifts decoupled from higher-frequency physiological noise.

The integration of EEG and fNIRS for depressive trait prediction reached a performance ceiling that underscores the strength of multimodal fusion in pBCI systems. In the optimal configuration, the system achieved:

- **Balanced Accuracy:** Central tendencies (mean and median) consistently surpassed the 90% threshold.
- **F1-Score:** Peak performance reached 0.867, indicating a high degree of precision and sensitivity in identifying sub-clinical depressive tendencies.

These metrics suggest that the “silent observer” approach, powered by end-to-end deep learning, can successfully bridge the gap between spontaneous neural dynamics and psychological screening, providing a high-accuracy alternative to active clinical assessments.

## IV. Conclusions

This study establishes the viability of a multimodal passive brain-computer interface (pBCI) for the objective identification of depressive traits through the end-to-end decoding of spontaneous neural dynamics. By implementing the Sinc-ShallowNet architecture, the system successfully bridges the gap between raw electro-hemodynamic signals and psychological screening, effectively bypassing the subjectivity and cognitive bias inherent in traditional self-report assessments. The findings confirm that the integration of EEG and fNIRS provides a uniquely robust physiological profile, wherein the high temporal resolution of electrical oscillations complements the spatial specificity of hemodynamic trends. This synergy reached its peak efficacy during auditory affective speech tasks, suggesting that the pBCI is particularly sensitive to the neural modulations triggered by verbal emotional stimuli.

The performance benchmarks achieved in this study validate the system’s high reliability for sub-clinical screening, with balanced accuracy consistently exceeding the 90% threshold and significantly outperforming the 50% chance-level baseline. Beyond mere classification, these results demonstrate that the proposed framework can isolate “silent” biomarkers within the 0.15 Hz hemodynamic range, providing a precise biological metric for early-stage intervention. Such objective screening mechanisms are essential for differentiating early depressive indicators from comorbid conditions, such as the initial stages of dementia in aging populations, or for identifying latent vulnerability in younger cohorts. Moving forward, the validation of this pBCI framework will be expanded from sub-clinical student populations to diverse clinical cohorts to further refine the longitudinal stability and diagnostic sensitivity of the system as a practical tool for data-driven psychiatric care.

## Acknowledgment

The authors express their gratitude to the student participants for their cooperation. Funding for this research was provided by the RIKEN Center for Advanced Intelligence Project (AIP) and BCI-Lab internal research funds.

